# Mobile Footprinting: Linking Individual Distinctiveness in Mobility Patterns to Mood, Sleep, and Brain Functional Connectivity

**DOI:** 10.1101/2021.05.17.444568

**Authors:** Cedric Huchuan Xia, Ian Barnett, Tinashe M. Tapera, Zaixu Cui, Tyler M. Moore, Azeez Adebimpe, Sage Rush-Goebel, Kayla Piiwaa, Kristin Murtha, Sophia Linguiti, Ellen Leibenluft, Melissa A. Brotman, Melissa Lynne Martin, Monica E. Calkins, David R. Roalf, Daniel H. Wolf, Danielle S. Bassett, David M. Lydon-Staley, Justin T. Baker, Lyle Ungar, Theodore D. Satterthwaite

**Affiliations:** Penn Lifespan Informatics and Neuroimaging Center, Department of Psychiatry, Perelman School of Medicine, University of Pennsylvania, Philadelphia, PA 19104, USA; Penn/CHOP Lifespan Brain Institute, University of Pennsylvania, Philadelphia, PA 19104, USA; Department of Biostatistics, Epidemiology and Informatics, Perelman School of Medicine, University of Pennsylvania, Philadelphia, PA 19104, USA; National Institute of Mental Health, Intramural Research Program, Bethesda, MD 20892; Department of Bioengineering, School of Engineering and Applied Science, University of Pennsylvania, PA 19104, USA; Department of Physics and Astronomy, University of Pennsylvania, PA 19104, USA; Department of Electrical & Systems Engineering, University of Pennsylvania, PA 19104, USA; Department of Neurology, Perelman School of Medicine, University of Pennsylvania, Philadelphia, PA 19104, USA; Santa Fe Institute, Santa Fe, NM 87501, USA; Annenberg School of Communication, University of Pennsylvania, Philadelphia, PA 19104, USA; Department of Psychiatry, Harvard Medical School, Boston, MA 02115; McLean Institute for Technology in Psychiatry, McLean Hospital, Belmont, MA 02478; Department of Computer and Information Science, School of Engineering and Applied Science, University of Pennsylvania, PA 19104, USA; Department of Genomics and Computational Biology, Perelman School of Medicine, University of Pennsylvania, Philadelphia, PA 19104, USA; Department of Operations, Information and Decisions, Wharton School, Philadelphia, PA 19104, USA; Department of Psychology, School of Arts and Sciences, Philadelphia, PA 19104, USA; Center for Biomedical Image Computation and Analytics, University of Pennsylvania, Philadelphia, PA 19104, USA; Penn Statistics in Imaging and Visualization Center, Department of Biostatistics, Epidemiology and Informatics, University of Pennsylvania, Philadelphia, PA 19104, USA; Leonard Davis Institute for Health Economics, University of Pennsylvania, PA 19104, USA

**Keywords:** smartphone, GPS, mobile phenotyping, fMRI, mood, affective instability

## Abstract

Mapping individual differences in behavior is fundamental to personalized neuroscience. Here, we establish that statistical patterns of smartphone-based mobility features represent unique “footprints” that allow individual identification. Critically, mobility footprints exhibit varying levels of person-specific distinctiveness and are associated with individual differences in affective instability, circadian irregularity, and brain functional connectivity. Together, this work suggests that real-world mobility patterns may provide an individual-specific signature linking brain, behavior, and mood.

## MAIN

Linking individual differences in behavior to brain function is a central task of behavioral neuroscience^1^. However, quantifying complex human behavior in real world settings remains a challenge. One alternative to standard behavioral assessment is *digital phenotyping*, which uses mobility data from personal smartphones to quantify moment-by-moment human behavior^2^. Prior work has associated geolocation features to important clinical outcomes in psychiatric disorders such as bipolar disorder and schizophrenia^3^, and has linked accelerometer metrics to post-surgical recovery^4,5^. Furthermore, researchers have recently begun to capitalize on the substantial variability of behavior assessed with digital phenotyping to link individual differences in brain and behavior. For example, lower prefrontal activity during processing negative emotions has been associated with individual exposure to urban green space^6^, while greater functional coupling of the hippocampus and striatum has been linked to location variability^7^.

While these studies suggest that digital phenotyping can be a powerful tool for studying individual differences, it remains unknown whether mobility patterns are in fact *person-specific*. Recent high-impact work has established that individual humans have unique patterns of functional brain connectivity^8,9^. The uniqueness of such brain-based “*fingerprints*” (also called “connectotypes*”*^10^) have been associated with development, cognition, and psychiatric conditions^11^. Establishing analogous person-specific mobility patterns – or mobility “*footprints*” – would constitute an important advance in behavioral neuroscience, and provide the foundation for targeted, individual-specific interventions. Accordingly, here we test the hypothesis that mobility patterns derived from personal smartphones can be used to create person-specific behavioral footprints. Furthermore, we evaluate whether the distinctiveness of these footprints was related to individual differences in mood, sleep, and brain functional connectivity.

As part of a study of trans-diagnostic affective instability in youth, we tracked 3,317 person-days of geolocation and 2,972 person-days of accelerometer data from 41 adolescents and young adults (28 females; mean [s.d.] age = 23.4 [3.5] years, range 17–30 years) – approximately 3 months per individual (**Fig. 1a**, **Supplementary Fig. 1**). In this sample, 93% of participants reported clinically significant affective instability in the context of psychiatric disorders (especially borderline personality disorder; see **Supplementary Table 1**). After applying hotdeck imputation to missing GPS data as implemented in the Smartphone Sensor Pipeline^13^, we constructed the daily mobility trajectory for each participant (**Fig. 1b** & **c**; see Online Methods). Instead of using raw coordinates that would allow trivial individual identification (and raise privacy concerns) given a participant’s exact location, we extracted high-level summary statistics of mobility features. These features (15 geolocation-based and 7 accelerometer-based) included time spent at home, number of locations visited, and many others (**Fig. 1d, Supplementary Table 2**).

**Figure 1.**
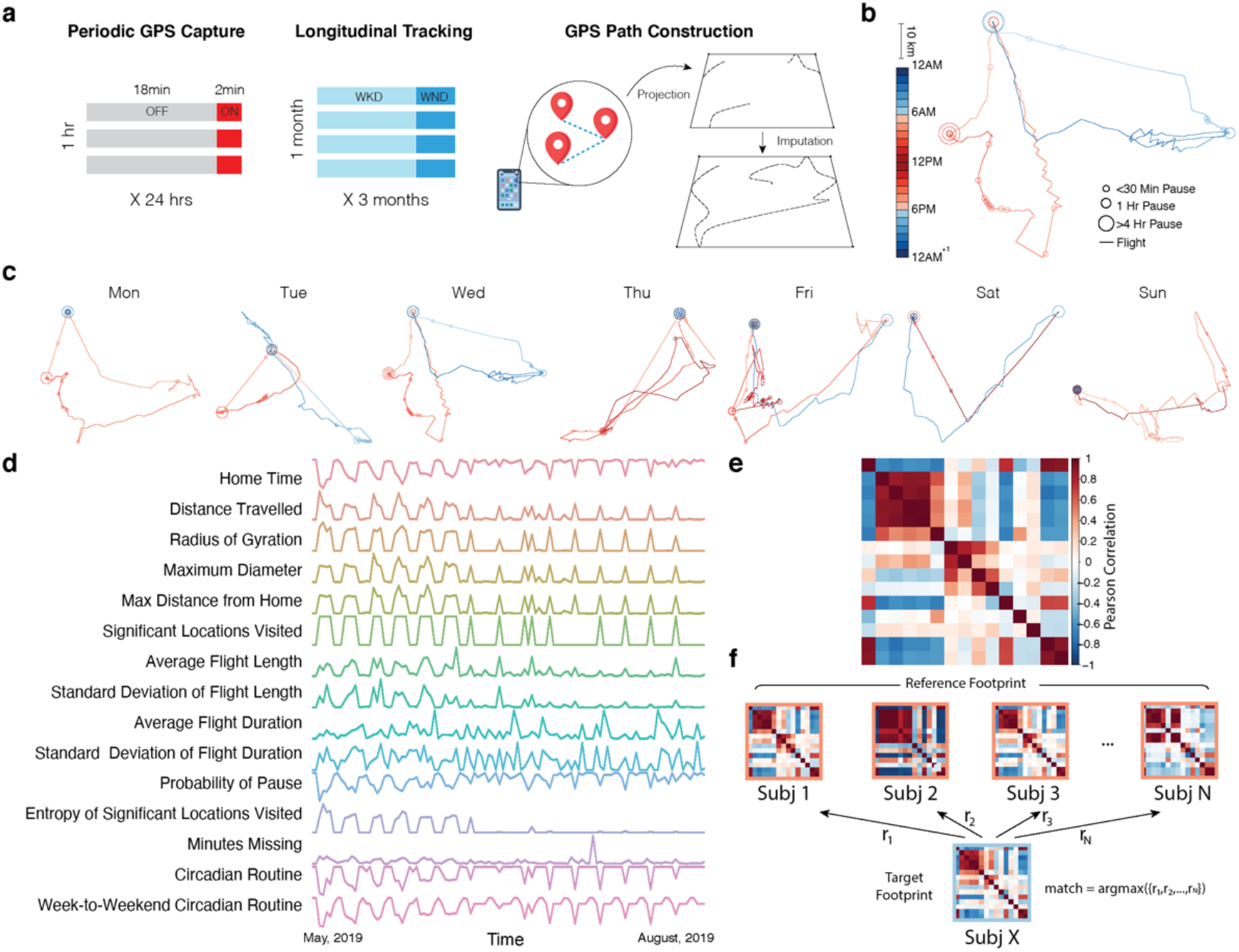
Constructing personal mobility “footprints”. **a**) We collected 3,317 person-days of mobility sensing data via personal smartphones from 41 adolescents and young adults. Geolocation data were recorded in cycles of 2min on and 18min off. Raw geolocation coordinates were de-identified via sphereto-2D standard space projection and were further imputed for missing data. **b**) For each individual, we constructed daily personal mobility trajectories, which consist of flights (movement) and pauses (stationary segments). Length of linear lines represents the duration of flights and size of circles represents the duration of pauses. Warm and cold colors indicate daytime and nighttime, respectively. **c**) A representative week of trajectories is shown, which demonstrates rich characteristics of personal mobility patterns formed over time. **d**) We extracted timeseries of mobility statistics (e.g. daily time spent at home) from geolocation and accelerometer data that parameterize movement characteristics over weeks to months. The example represented all 110 days of participants’ geolocation metrics recorded. **e**) For each individual, we constructed a covariance matrix from the mobility metric timeseries. Each cell of the matrix was populated by the Pearson correlation between a given pair of mobility metrics. Warm and cold colors indicate positive and negative correlations, respectively. **f**) We randomly divided data into two equally sized parts, called the reference and target set. *Subj X* from the target set was matched to the subject in the reference that had the highest correlations between their footprints (*argmax(r_1_, r_2_, ..r_N_)*). The identification was considered correct when underlying data came from the same subject; otherwise, the identification was considered incorrect. We quantified individual identification accuracy as the proportion of correct identifications across the entire sample; this procedure was repeated 1,000 times across different random partitions of the data.

When tracked over weeks to months, these timeseries of mobility statistics captured rich characteristics of individual mobility patterns. One illustrative example of the sensitivity of the timeseries to track mobility patterns is when COVID-19 pandemic hit the Philadelphia area towards the end of the study period. Participants who were still engaged in active data collection (n=3) exhibited dramatic shifts in mobility features (**Supplementary Fig. 2**). Of note, as the data points during COVID-19 represented merely 1.1% of all data, the findings reported below did not change significantly when these data were removed.

Drawing on prior work of brain connectome “fingerprinting,”^8,10^we created a covariance matrix of each participant’s fifteen geolocation-based and seven accelerometer-based mobility features timeseries to identify individuals (**Fig. 1e**), akin to a person-specific mobility “footprint.” Data from each individual is was partitioned into two groups: the target partition and the reference partition. For each individual, the data in the target partition was separately correlated with every individual’s data in the reference partition; this procedure yielded 41 correlation values. A correct identification was declared only when the maximum correlation was from the data belonging to the same individual across the target and reference partitions **(Fig. 1f)**. In order to ensure that the random partitioning of the data did not impact results, this matching procedure was then repeated for each individual 1000 times (Online Methods).

Initial inspection across random partitions of the data revealed that it was visually apparent that there was substantially greater correlation between mobility footprints *within* participants rather than *between* participants (**Fig. 2a**). Permutation testing on the entire sample revealed that individuals could be successfully identified using their mobility footprints (*p* <0.001; **Fig. 2b**). Across 1,000 random data partitions, the mean individual identification accuracy was 63%. Critically, this accuracy was far better than chance performance determined by a permuted null distribution (mean: 3% accuracy; see **Fig 2b** inset).

**Figure 2.**
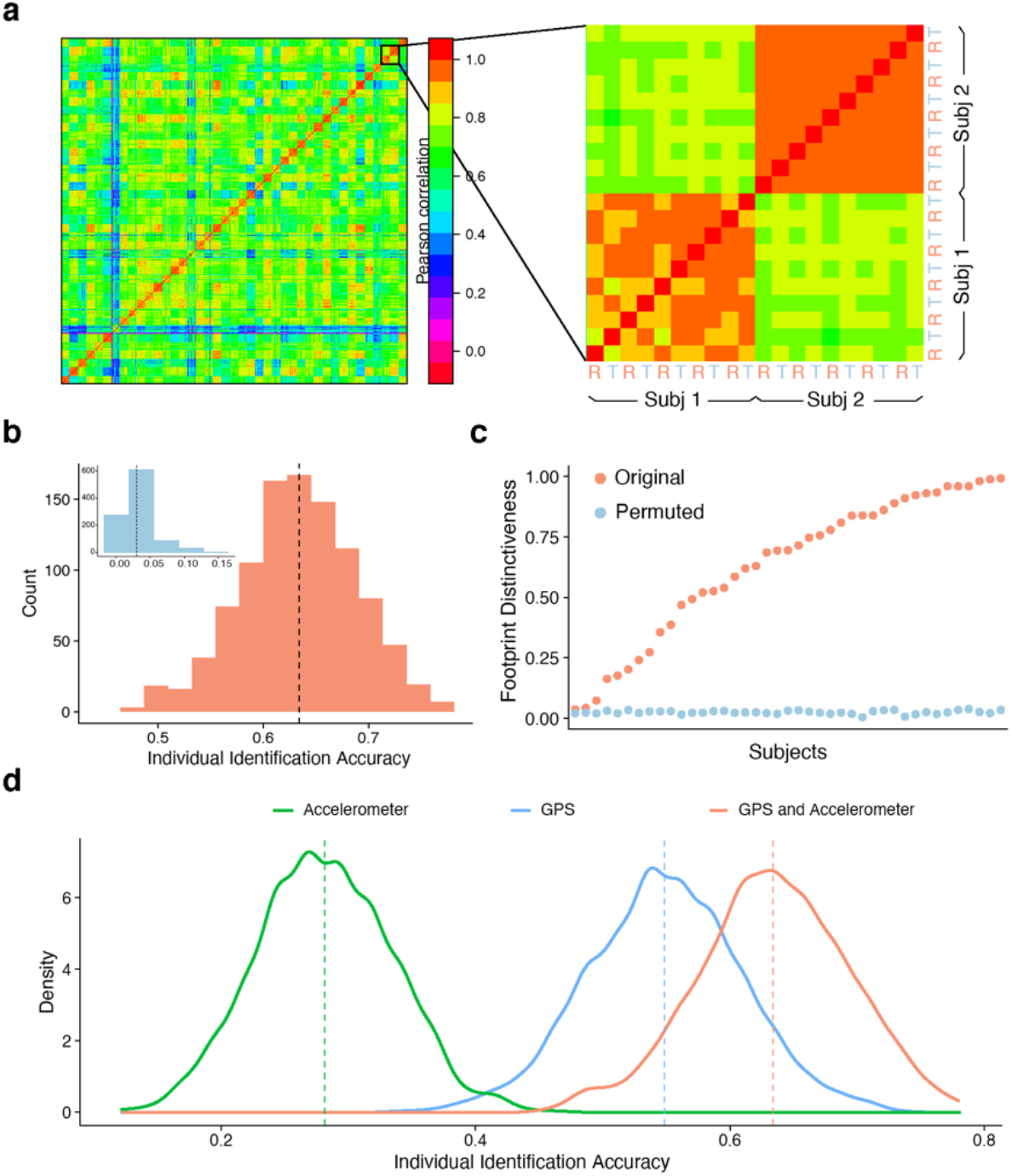
Identifying individuals using personal footprints. **a**) As an initial step, we visualized the similarity of mobility features across multiple random reference and target partitions (R & T in inset). It was readily apparent that mobility features were more highly correlated within participants (on diagonal) across data partitions than between participants (off diagonal). Note that this visualization was not used in statistical analysis or individual identification. **b**) Across 1,000 random partitions, mobility footprinting enabled successful individual identification (mean: 63%, S.D.: 6%). In contrast, the mean chance accuracy from 1,000 permutation was 3% (inset, *p* <0.001). **c**) For each individual, we calculated the footprint distinctiveness, or the percentage of correct identification across the 1,000 random partitions of the data. Ranked in ascending order, participants’ footprint distinctiveness exhibited a wide range, from 4% to 99%. However, even the participant with the lowest identification distinctiveness was significantly higher than the null distribution. **d**) Individual identification based on geolocation alone had higher accuracy than accelerometer alone. However, they appeared to encode complementary features, as performance was maximal when both measures were used in footprinting.

Moving beyond aggregate measures of accuracy across the group, we next investigated whether certain individuals could be consistently identified more accurately than others. Similar to prior studies of brain connectome fingerprinting^8,10,11^, we refer to this measure as an individual’s “*footprint distinctiveness*”. Notably, individuals exhibited a wide distribution of footprint distinctiveness, ranging from 4% to 99% (**Fig. 2c**). In other words, certain participants had such distinct mobility patterns that it enabled correct identification nearly every single time; other participants were difficult to identify. Nonetheless, permutation testing showed that all participants had significant footprint distinctiveness compared to the null distribution.

As the group and individual level accuracy results reported thus far were based on the combination of geolocation and accelerometer features, we next examined each feature set separately. Individual footprint distinctiveness derived by geolocation was not correlated with that of accelerometer (*r* = 0.18; *p* = 0.26). Interestingly, while accelerometer data alone yielded lower identification accuracy (28%) than geolocation data (55%), combining these features resulted in higher identification accuracy, suggesting that they encode complementary information (**Fig. 2d**). Importantly, individual identification accuracy was stable across different inclusion thresholds for data missingness and was robust to removal of individual mobility features (**Supplementary Fig. 3**).

We next investigated participant factors that influenced footprint distinctiveness. We found that data quantity (i.e. number of days recorded) was associated with footprint distinctiveness (**Supplementary Fig. 4**). In contrast, the amount of missing data *within* a given day was unrelated to footprint distinctiveness. Based on this result, all subsequent analyses of individual differences related of footprint distinctiveness controlled for number of days of data available. As a next step, we evaluated whether footprint distinctiveness was related to age or sex in our sample of adolescents and young adults. We found that geolocation-based footprints became more distinct with age across the transition from adolescence to adulthood (partial *r* = 0.33, *p* <0.05, **Supplementary Fig. 5**). Furthermore, female sex was associated with higher accelerometer-based footprint distinctiveness (*Cohen’s d* = 1.27, *p* < 0.001, **Supplementary Fig. 5**).

We next evaluated how footprint distinctiveness was related to a key domain of psychopathology: affective instability. Affective instability is a major feature of many psychiatric disorders^14^, including borderline personality disorder. Affective instability is particularly prominent in youth^15^, and is an important predictor of suicide^16^. However, affective instability is often challenging to quantify using standard tools as it is fundamentally a dynamic measure^17^. We capitalized on participant ratings of multiple mood features collected three times a day for two weeks using ecological momentary assessment in order to quantify affective instability. We hypothesized that individuals who had less predictable patterns of mobility (i.e., reduced footprint distinctiveness) would have higher levels of affective instability. While controlling for data quantity, age, sex, and the mean of mood ratings, we found that affective instability (measured by root mean square of successive differences^18^) was associated with reduced footprint distinctiveness (partial *r* = −0.37, *p* < 0.05, **Fig. 3a**). Furthermore, given well-established links between sleep disturbance and mood disorders^19^, we also evaluated whether variability in sleep duration was also associated with footprint distinctiveness. While controlling for covariates as above, we found that variability in sleep duration was similarly associated with reduced footprint distinctiveness (partial *r* = −0.36, *p* < 0.05, **Fig. 3b**).

**Figure 3.**
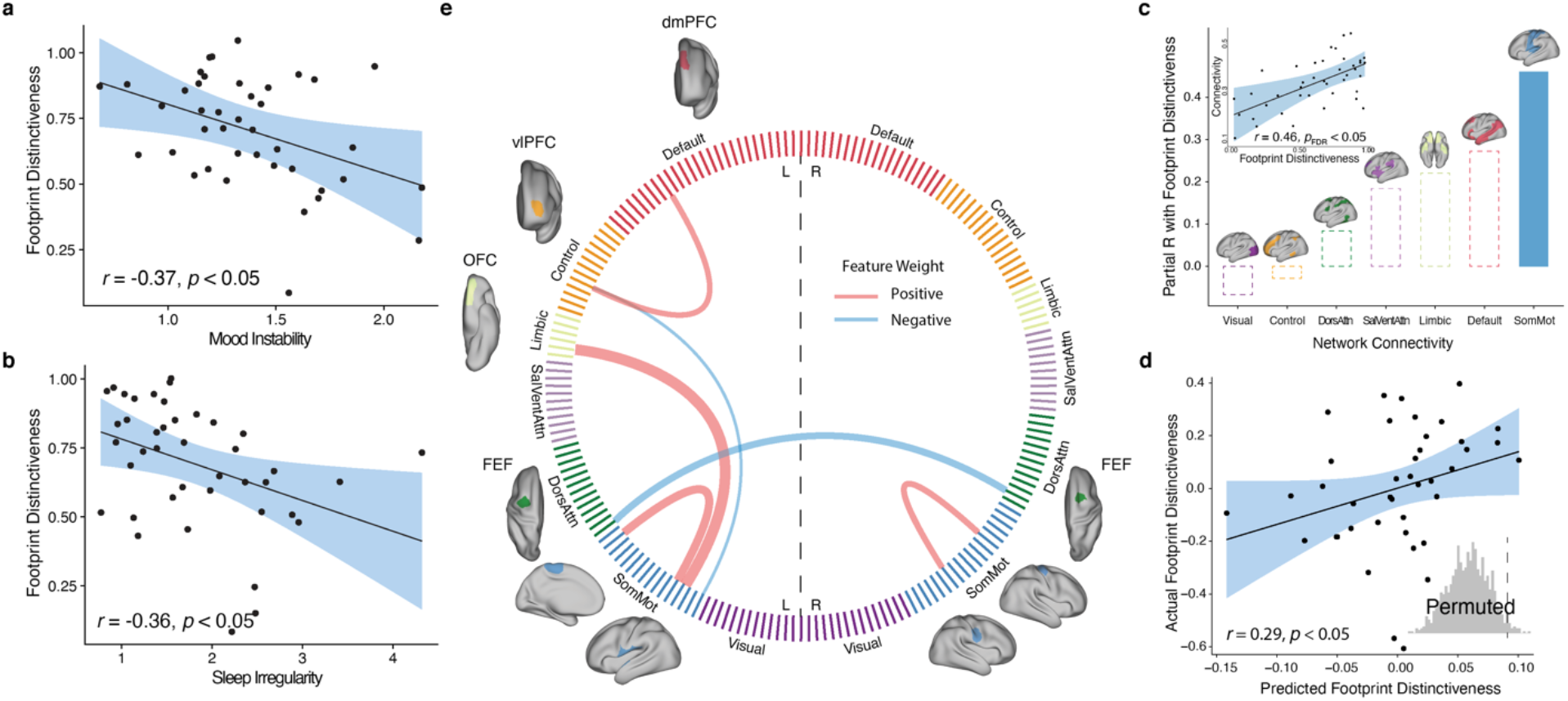
Individual footprint distinctiveness is associated with affective instability, sleep irregularity, and patterns of brain functional connectivity. **a**) Greater affective instability, measured by root mean square of successive differences in mood measures from ecological momentary assessment items acquired three times a day, was associated with reduced footprint distinctiveness (*r* = −0.37, *p* < 0.05), after controlling for data quantity, age, sex, and mean level of mood ratings. **b**) Similarly, we found that increased variability in sleep duration was associated with reduced footprint distinctiveness (*r* = −0.36, *p* < 0.05), after controlling for covariates. **c**) Across functional brain networks, only greater connectivity within the somatomotor network had a significant association with footprint distinctiveness (*r* = 0.46, *p* < 0.05, corrected for multiple comparisons with the false discovery rate). **d**) Patterns of brain functional connectivity significantly predicted individual footprint distinctiveness using leave-one-out cross-validation (*r* = 0.29, inset: permutation-based *p* = 0.025). **e**) Six network edges consistently contributed to the sparse regression model. These edges included greater connectivity within somatomotor network, reduced connectivity between left and right frontal eye fields (FEF), increased connectivity between the somatomotor network and the left orbital frontal cortex (OFC) in the limbic network, as well as increased connectivity between the vlPFC (ventrolateral prefrontal cortex in the frontoparietal network) and the dmPFC (dorsolateral prefrontal cortex in the default mode network). Cord thickness reflects the weights in the model, reflecting each edge’s contribution to the prediction; cord color indicates the sign of the weights.

As a final step, we investigated whether footprint distinctiveness was related to patterns of functional connectivity. Initially, we examined associations with a simple summary measure of high-dimensional functional connectivity data: the mean connectivity within each of seven canonical large-scale functional networks^20^. While controlling for covariates as above (as well as in-scanner motion) and correcting for multiple comparisons with the false discovery rate, we found that footprint distinctiveness was associated with greater connectivity within the somatomotor network (*r* = 0.46, *p*_fdr_ = 0.03, **Fig. 3c**). Previous work has demonstrated that somatomotor network connectivity develops over the lifespan (years)^21^and is altered acutely (days) during limb disuse^22^; our results further suggest that mobility patterns over weeks-months can be predicted by somatomotor network connectivity.

Lastly, we moved beyond the simple summary measure of mean network connectivity and investigated whether complex multivariate patterns of functional connectivity could predict footprint distinctiveness in unseen data. Given that there were far larger number of features than participants, we used regularized regression with leave-one-out cross-validation and nested parameter tuning, followed by permutation testing to determine significance (Online Methods). We found that multivariate patterns of functional connectivity could predict footprint distinctiveness in unseen data (*r* = 0.29, *p* = 0.025; **Fig. 3d**). The predictive model yielded results that aligned with the mass-univariate analyses (**Fig. 3c**), suggesting that the multivariate model was driven in part by features linked to somatomotor network (67% of edges selected by the model). Moreover, this model also revealed important features beyond the motor system, including increased connectivity between the frontoparietal and default mode system (**Fig. 3e**).

Taken together, these results establish that mobility patterns collected from smartphones can be used to create a person-specific footprint. Notably, the distinctiveness of this footprint increased with age, was reduced in association with both affective instability and circadian irregularity, and was related to patterns of functional brain connectivity. These results align with impactful prior work on “connectome fingerprinting”^8^(or “connectotyping”^10^), which have shown that individuals can be identified based on their pattern of functional connectivity. Interestingly, result from these prior studies have shown that – like the footprint distinctiveness examined here – connectome distinctiveness increases with age and is reduced in association with psychiatric symptoms^11^.

Our finding that footprint distinctiveness is related to data quantity recalls recent work demonstrating that the ability to delineate person-specific functional brain networks is dependent in large part on the quantity of data available^23,24^. However, while accruing large amounts of functional imaging data is often difficult and expensive, passive collection of long timeseries of mobility data is both tolerable for participants and inexpensive. The high degree of scalability enabled by ubiquitous usage of personal smartphones will allow future studies to test the generalizability of these findings across different age groups and clinical samples. Moving forward, mobility-based digital biomarkers that combine objective measurement and individual-specific analysis of behavior may accelerate the advances in personalized diagnostics for diverse psychiatric illnesses.

## ONLINE METHODS

### Participants

A sample of 41 adolescents and young adults (28 females; mean (s.d.) age = 23.4 (3.5) years, range 17–30 years) were enrolled as part of a study of affective instability in youth. Participants were recruited via the Penn/CHOP Lifespan Brain Institute or through the Outpatient Psychiatry Clinic at the University of Pennsylvania. Of these 41 participants, 38 participants met criteria for Axis I psychiatric diagnosis based on a semi-structured clinical interview^1^; 33 met criteria for more than one disorder (**Supplementary Table 1**). Additionally, 16 of the 41 participants met criteria for a personality disorder (mainly borderline personality disorder) based on assessment with the SCID-II^1^. All participants provided informed consent to all study procedures; for minors, the parent or guardians provided informed consent and the minor assented as well. This study was approved by the University of Pennsylvania Institutional Review Board.

### Mobility data acquisition

Global Positioning System (GPS) geolocation data were acquired via the Beiwe platform^2^. Participants were asked to download the Beiwe application on their personal smartphone. The application recorded the location of the participant’s phone in latitude, longitude, and altitude, as well as the precision of that measure. To conserve battery and minimize degradation of the phone performance, Beiwe was designed to track participant’s geolocation in a periodic fashion. Specifically, Beiwe tracked GPS for 2 minutes every 20 minutes, resulting in 144 minutes of data recording and 1296 minutes of dormancy in a 24-hour cycle. Due to user and device related factors in the naturalistic setting, such as phone powered off, no cell signal, or airplane mode, longer periods of recording dormancy were possible. Mobility data were automatically uploaded via WiFi to a cloud-based data management system daily.

In total, 3,317 days of GPS tracking across all participants were obtained (mean (s.d.) = 77 (26) days, range 14–132 days, see **Supplementary Figure 1**). After removing the first and last days of each participant’s study period when only partial data were recorded and days containing no data, the remaining data available for analysis had 3,156 days.

Accelerometer data were also acquired via the Beiwe platform. The application recorded the participants’ acceleration in three cardinal axes (x, y, and z) in m/s^2^. In total, 2,972 days of accelerometer data were obtained across all participants (mean (s.d.) = 74 (32) days, range 15-134 days). After removing the first and last days of each participant’s study period when only partial data were recorded, the remaining data available for analysis had 2972 days.

### Mobility data processing

#### GPS data preprocessing

Raw GPS data were processed using the Smartphone Sensor Pipeline^3^, a validated pipeline specifically designed to handle GPS data while accounting for data missingness. First, each subject’s GPS longitude and latitude coordinates on the spherical Earth’s surface were transformed to a standardized two-dimensional Cartesian plane, thus deidentifying subject’s realworld locations. Second, the data were converted to a sequence of flights and pauses, where flights were defined as segments of linear movements and pauses were defined as periods of no movement. Finally, missing flights and pauses were then imputed by the hot-deck method^4^, which resamples from observed events over each missing interval.

#### Mobility metrics calculation

Using the constructed subject mobility traces and the Smartphone Sensor Pipeline, 15 GPS-based mobility metrics were calculated for each day of recording, defined as midnight to midnight. See Barnett et al. for details^3^. An additional seven accelerometer-based mobility metrics were calculated for each day of recording. These were implemented according to methods described in the RAPIDS pipeline^5^. See **Supplementary Table 2** for definitions of each metric.

#### Mobility footprint construction

Inspired by person-specific connectome fingerprints^6,7^, we constructed a mobility footprint for each participant using the covariance matrix of mobility metrics. First, we extracted the mobility metric time series by concatenating the daily mobile metric output from the Smartphone Sensor Pipeline. Then we computed the pairwise Pearson correlation for all the mobility metrics to construct a covariance matrix. The nodes of the network were the mobility metrics, and the edges of the network were the Pearson correlation coefficients between metrics. We refer to the resulting covariance matrix as the “Mobility Footprint.” This procedure was carried out separately for GPS- and accelerometer-based mobility data. For the main analysis, the upper triangle of the resulting covariance matrices from GPS and accelerometer metrics were concatenated and were used as input features for the individual identification procedure. We also repeated the identification procedure using GPS or accelerometer features alone.

As a sensitivity analysis to test performance of alternative features for individual identification, we also computed the mean and the stability of each measure and used these features to identify participants. Stability was defined as the root mean square of the successive differences (RMSSD)^8^of each measure (**Supplementary Figure 6**).

#### Individual identification procedure (“footprinting”)

We randomly partitioned each subject’s data into two equally sized parts, named the “reference” and the “target”, respectively^6^. The objective of the individual identification procedure was to match the subject from the target group to the same one in the reference group. For a given subject, *S*, we computed the Pearson correlation (*r*) between that subject’s features in the target group, *S_T_*, and everyone’s features in the reference group, *S_R_ 1, S_R_2, S_R_N*, where *N* is the total number of participants.

Individual identification was operationalized as the maximum of the resulting *r_1_, r_2_, … r_3_*. In other words, when the subject in the reference group having the mobility features that maximally correlated with that of the target subject, these two participants were declared correctly matched:

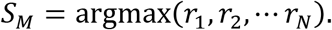

The individual identification accuracy was the number of correct identifications divided by the total number of random data partitions *P*:

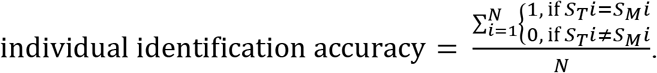

The above individual identification procedure was repeated 1,000 times, each time with a new random data partition (P). We calculated the average individual identification accuracy across the 1,000 runs, which yielded a distribution of sample-wise identification accuracy. Furthermore, we also calculated the accuracy for each participant, defined as the number of correct identifications for that specific participant divided by the number of data partitions (*B*). We refer to this participant-specific identification accuracy as the individual footprint distinctiveness:

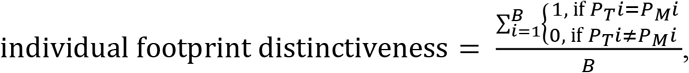

where *P_T_ i* is target in a partition for subject *i*, and *P_M_ i* is matched subject. We conducted the individual identification procedure using the covariance matrix of the GPS data, accelerometer data, as well as the combined feature set. Sensitivity analyses examined the mean and variance of each feature.

#### Similarity matrix construction

To visualize the individual footprint distinctiveness, we constructed similarity matrices between participants’ mobility covariance features^9^. First, we concatenated the daily mobility metrics for a participant from multiple random data partitions. Next, a similarity matrix was constructed by computing the Pearson correlation coefficients between every pair of participants. The resulting matrix was a symmetric matrix, where the nodes were each participant and the edges were the correlation coefficients between any two participant’s mobility metrics. This grouping procedure was performed solely for visualization, highlighting the within-individual, across-partition block structures on the diagonal of the matrix. This grouping was not used in any statistical analysis.

#### Permutation testing

To assess the statistical significance of individual identification accuracy, we used a permutation testing procedure to create a null distribution of accuracy. Specifically, we randomly scrambled the identity of the daily mobility metrics, thus disrupting the linkage between the mobility data and the corresponding participant. We repeated the individual identification procedure for each random permutation. The empirical *p-*value was then calculated as the proportion of times when the permuted data yielded higher accuracy than the original data:

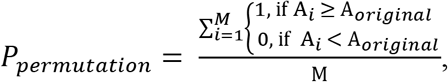

where *A* is the individual identification accuracy, and *M* is the total permutations.

#### Sensitivity analysis of data missingness

To understand the effect of data missingness on our ability to identify participants’ mobility footprint, we conducted sensitivity analyses that used four sets of data constructed using different thresholds for data missingness^3,10^. Specifically, we applied four thresholds with diminishing tolerance for the number of missing samples (i.e., minutes recorded) in a day’s worth of data to be included in analysis (**Supplementary Figure 1**). At the 100^th^ percentile level, which corresponded to retaining all available days except for those with all data missing (or 1,440 minutes), 79 recording days were removed, which resulted 3,156 days remaining for analysis. At the 90^th^ percentile, a further 216 days were removed, yielding 2,940 days for analysis. At 80^th^ percentile, a further 356 days were removed, resulting in 2,584 days for analysis. Finally, at 75^th^ percentile, a further 171 days were removed, resulting in 2,413 days remaining for analysis. Using these four sub-samples constructed with different inclusion criteria, we then repeated the individual identification procedure and permutation testing as described above.

#### Feature lesion analysis

To further investigate the influence of any single feature’s influence on the individual identification accuracy, we conducted a feature lesion analysis. We sequentially removed one metric (out of the total 15 geolocation mobility metrics available) and constructed a new covariance matrix which had one node (and 14 edges) less than the original feature covariance matrix. Using this reduced feature set, we repeated the individual identification and permutation testing procedures as described above (**Supplementary Figure 3**).

#### Ecological momentary assessment

Using the Beiwe platform application on personal smartphones, participants completed daily questionnaires specifically designed to assess mood variability at three timepoints throughout the day^11^. In each survey, participants rated on a scale from 1 (“not at all”) to 7 (“extremely”) of their endorsement of seven statements assessing mood variability, aggression, impulsivity, and self-esteem since the last time they had answered the survey to capture their mood (**Supplementary Table 3**). All seven items were concatenated to create an overall mood scale. Additionally, every morning, participants were also asked about their sleep patterns and quality from the night before. To quantify the variability of answers to the mood survey, we calculated the root mean square of successive differences (RMSSD) between concatenated answers. Similarly, we also calculated the RMSSD of sleep duration as a measurement of its stability. We built a generalized additive model (GAM) to investigate the association between mood and sleep duration stability while accounting for covariate effects including data quantity, sex, age, and mean levels of the measure. Age was modeled using penalized splines within GAM using restricted maximum likelihood (REML) to estimate linear and nonlinear developmental effects without over-fitting the data^12,13^.

### Functional Connectivity Analysis

#### Imaging Acquisition

As previously described^14^, structural and functional MRI scans were acquired using in a single session on a clinically-approved 3 Tesla Siemens Prisma (Erlangen, Germany) quadrature body-coil scanner and a Siemens receive-only 64-channel head coil at the Hospital of the University of Pennsylvania. Prior to functional MRI acquisitions, a 5-min magnetization-prepared, rapid acquisition gradient-echo T1-weighted (MPRAGE) image (TR = 1810 ms; TE = 3.45 ms; TI = 1100 ms, FOV = 180 × 240 mm^2^, matrix = 192 × 256, 160 slices, effective voxel resolution = 0.9375 × 0.9375 × 1 mm^3^) was acquired. We used one resting-state (1200 volumes) scan as part of this study. All fMRI images were acquired with the same multi-band, interleaved multi-slice, gradient-echo, echo planar imaging (GE-EPI) sequence sensitive to BOLD contrast with the following parameters: TR = 500 ms; TE = 25 ms; multiband acceleration factor = 6, flip angle = 30°; FOV = 192 × 192 mm^2^; matrix = 64 × 64; 48 slices; slice thickness/gap = 3/0 mm, effective voxel resolution = 3.0 × 3.0 × 3.0 mm^3^.

#### Image Processing

All preprocessing was performed using fMRIPrep 20.0.7^15^, which is based on Nipype 1.4.2^16^, and XCP Engine^17,18^(PennBBL/xcpEngine: atlas in MNI2009 Version 1.2.3; Zenodo: http://doi.org/10.5281/zenodo.4010846). The T1-weighted (T1w) image was corrected for intensity non-uniformity (INU) with N4BiasFieldCorrection^19^, distributed with ANTs 2.2.0^20^, and used as T1w-reference throughout the workflow. The T1w-reference was then skull-stripped with a Nipype implementation of the antsBrainExtraction.sh workflow (from ANTs), using OASIS30ANTs as target template. Brain tissue segmentation of cerebrospinal fluid (CSF), white-matter (WM) and gray-matter (GM) was performed on the brain-extracted T1w using FAST in FSL 5.0.9^21^. Volume-based spatial normalization to MNI2009c standard space was performed through nonlinear registration with antsRegistration (ANTs 2.2.0), using brain extracted versions of both the T1w reference and the T1w template.

BOLD runs were first slice-time corrected using 3dTshift from AFNI 20160207^22^and then motion corrected using mcflirt (FSL 5.0.9)^21^. A fieldmap was estimated based on a phase difference map calculated with a dual-echo GRE sequence, processed with a custom workflow of SDCFlows inspired by the epidewarp.fsl script and further improvements in HCP Pipelines^23^. The fieldmap was then co-registered to the target EPI reference run and converted to a displacement field map with FSL’s fugue and other SDCflows tools. Based on the estimated susceptibility distortion, a corrected BOLD reference was calculated for a more accurate coregistration with the anatomical reference. The BOLD reference was then co-registered to the T1w reference using bbregister (FreeSurfer) which implements boundary-based registration^24^. Co-registration was configured with nine degrees of freedom to account for distortions remaining in the BOLD reference. Six head-motion parameters (corresponding rotation and translation parameters) were estimated before any spatiotemporal filtering using mcflirt. Finally, the motion correcting transformations, field distortion correcting warp, BOLD-to-T1w transformation and T1w-to-template (MNI) warp were concatenated and applied to the BOLD timeseries in a single step using antsApplyTransforms (ANTs) with Lanczos interpolation.

After pre-processing with fMRIPRep, confound regression was carried out in XCP Engine. Preprocessed timeseries were despiked and then de-noised using a 36-parameter confound regression model that has been shown to minimize the impact of motion artifact^25^. Specifically, the confound regression model included the six framewise estimates of motion, the mean signal extracted from eroded white matter and cerebrospinal fluid compartments, the global signal, the derivatives of each of these nine parameters, and quadratic terms of each of the nine parameters as well as their derivatives. Both the BOLD-weighted time series and the confound regressor timeseries were temporally filtered simultaneously using a fist-order Butterworth filter with a passband between 0.01 and 0.08 Hz to avoid mismatch in the temporal domain^26^. Confound regression was performed using AFNI’s 3dTproject. Note that in-scanner head motion was also included as a covariate in all regression models (see below).

#### Functional network and community connectivity

Functional connectivity between each pair of brain regions was quantified as the Fisher-transformed Pearson correlation coefficient between the mean regional BOLD time series. For each participant, a 200 × 200 weighted adjacency matrix encoding the connectome was constructed^27^. Each node was assigned to one of seven canonical functional brain modules or communities defined by Yeo et al^28^.

The within-community connectivity is defined as

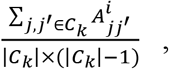

Where 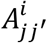 is the weighted edge strength between the node *j* and node *j*′, both of which belong to the same community *C_k_*, for the *i*-th subject. The cardinality of the community assignment vector, *C_k_*, represents the number of nodes in the *k*-th community^29^.

#### Mass-univariate analysis

For each of the seven canonical networks, we fit generalized additive model (GAM) to investigate the relationship between within-network connectivity and footprint distinctiveness, while controlling for in-scanner motion, mobility data quantity, sex, and age. Specifically, we used penalized splines using restricted maximum likelihood (REML) within GAM to estimate linear and nonlinear age-related changes^12,13^. We controlled for multiple comparisons using the False Discovery Rate (*Q*<0.05).

#### Predicting footprint distinctiveness using functional connectivity

We fit a penalized regression model to predict footprint distinctiveness using brain functional connectivity^6^. In each iteration of leave-one-out cross-validation, one subject was left out as the testing set and the rest the training set. Using the training set, we computed residualized footprint distinctiveness from a GAM model with covariates as above (linear terms for in-scanner motion, data quantity, sex; age was modeled with as a penalized spline). Then we fit a lasso regression model to predict the residualized footprint distinctiveness using a sparse collection of functional connectivity edges. L1 lasso hyperparameter was tuned in a nested leave-one-out fashion. Next, we calculated the predicted footprint distinctiveness for the unseen subject in the testing set. After all iterations, we obtained predicted footprint distinctiveness for all participants and then calculated the Pearson correlation between the actual footprint distinctiveness and predicted values.

## Code availability and data access

The code for GPS data preprocessing, mobility metric extraction, individual identification, additional analysis, and data visualization is available in R on github: https://github.com/PennLINC/footprinting

Code notebook is available at: https://pennlinc.github.io/footprinting/

Data is available upon request.

