## Supplementary Material for "Mobile Footprinting: Linking Individual Distinctiveness in Mobility Patterns to Mood, Sleep, and Brain Functional Connectivity"

**Supplementary Table 1 | Psychiatric diagnoses of participants**

| Axis I Diagnosis | | Axis II Diagnosis | |
|---|---|---|---|
| Total | n=41 | | n=41 |
| No Diagnosis | n=3 | No Diagnosis | n=7 |
| Diagnosis | n=38 | Borderline personality disorder | n=12 |
| Major depressive disorder | 24 | Personality disorder NOS | n=4 |
| Bipolar disorder | 4 | | |
| Depressive disorder NOS | 1 | | |
| Mood disorder NOS | 1 | | |
| Generalized anxiety disorder | 14 | | |
| Post-traumatic stress disorder | 14 | | |
| Social phobia | 12 | | |
| Obsessive-compulsive disorder | 11 | | |
| Panic disorder | 5 | | |
| Anxiety disorder NOS | 2 | | |
| Attention-Deficit Hyperactivity Disorder | 6 | | |
| Schizoaffective Disorder | 1 | | |
| Substance Related Disorders | 18 | | |

**Supplementary Table 2 | Footprint mobility metrics**

| | GPS Mobility Metric[1] | Definition | Comments |
|---|---|---|---|
| 1 | Home Time | Amount of time (minutes) spent within a 200-meter radius of home | Home location is determined by the location with the largest total amount of time between 9PM and 6AM over the course of the study period. |
| 2 | Distance Travelled | Sum of the flight lengths (meters) over all flights that occurred on that day | Flights are quantified as segments of linear movements. |
| 3 | Radius of Gyration | Root mean square distance (meters) from a central location | As radius of gyration quantifies the distribution around a central location, it acts as a proxy for the area covered in a day. |
| 4 | Maximum Diameter | Maximum pairwise distance (meters) between any two pause locations that both occur on the same day | |
| 5 | Maximum Distance from Home | Maximum of the distances (meters) between home and every other pause location. | |
| 6 | Significant Locations Visited | Number of significant locations a subject visited in a day | The set of significant locations is determined by a K-means procedure on all pause locations with greater than 10 minutes |
| 7 | Average Flight Length | Average length (meters) of all flights that occur on that day | |
| 8 | Standard Deviation of Flight Length | Standard deviation of flight length (meters) over all flights that occur on that day | |
| 9 | Average Flight Duration | Average duration (seconds) of all flights that occur on that day | |
| 10 | Standard Deviation of Flight Duration | Standard deviation of flight duration (seconds) over all flights that occur on that day | |
| 11 | Probability of Pauses | Fraction of time stationary relative to the amount of time spent moving | Pauses are defined as periods of no movement |
| 12 | Entropy of Significant Locations Visited | Randomness of the locations a subject visited in a day | Large values indicate a subject spreading their time out across many different locations fairly evenly for that day, whereas small values indicate a concentration at few significant locations |
| 13 | Minutes Missing | Number of minutes of missing data for that day | |
| 14 | Circadian Routine | Degree to which a participant followed their daily routine on a given day | Low values indicate a break from routine on a given day, whereas high values indicate that the subject followed their daily routine |
| 15 | Weekend/Weekday Circadian Routine | Degree to which a participant followed their weekend or weekday routine | |

| | Accelerometer Mobility Metric[2] | Definition | Comments |
|---|---|---|---|
| 1 | Maximum Magnitude | The maximum magnitude of acceleration, defined by $\|acceleration\| = \sqrt{x^2 + y^2 + z^2}$. | |
| 2 | Minimum Magnitude | The minimum magnitude of acceleration. | |
| 3 | Average Magnitude | The average magnitude of acceleration. | |
| 4 | Median Magnitude | The median magnitude of acceleration. | |
| 5 | Standard Deviation Magnitude | The standard deviation of acceleration. | |
| 6 | Absolute Duration of Exertional Activity | Total duration of daily exertional activity based on acceleration | The threshold for defining exertional vs. non-exertional activity was set at 0.15g[2], as described in Panda et al[3]. |
| 7 | Relative Duration of Exertional Activity | Duration of daily exertional activity relative to non-exertional activity | |

**Supplementary Table 3 | Ecological Momentary Assessment of Mood**

| | Since last beep…, |
|---|---|
| **1** | my mood has changed a lot.<br>1 = not at all; 7 = extremely |
| **2** | I have felt really, really angry or out of control.<br>1 = not at all; 7 = extremely |
| **3** | I thought I am no good, a bad person, or that nobody loves me.<br>1 = not at all; 7 = extremely |
| **4** | I have been social versus isolated myself from others.<br>1 = extremely isolated; 7 = very social |
| **5** | I had an argument.<br>1 = yes; 2 = no |
| **6** | my social interactions have been overall positive versus negative.<br>1 = very positive; 7 = very negative |
| **7** | I have done something impulsive or risky.<br>1 = yes; 2 = no |

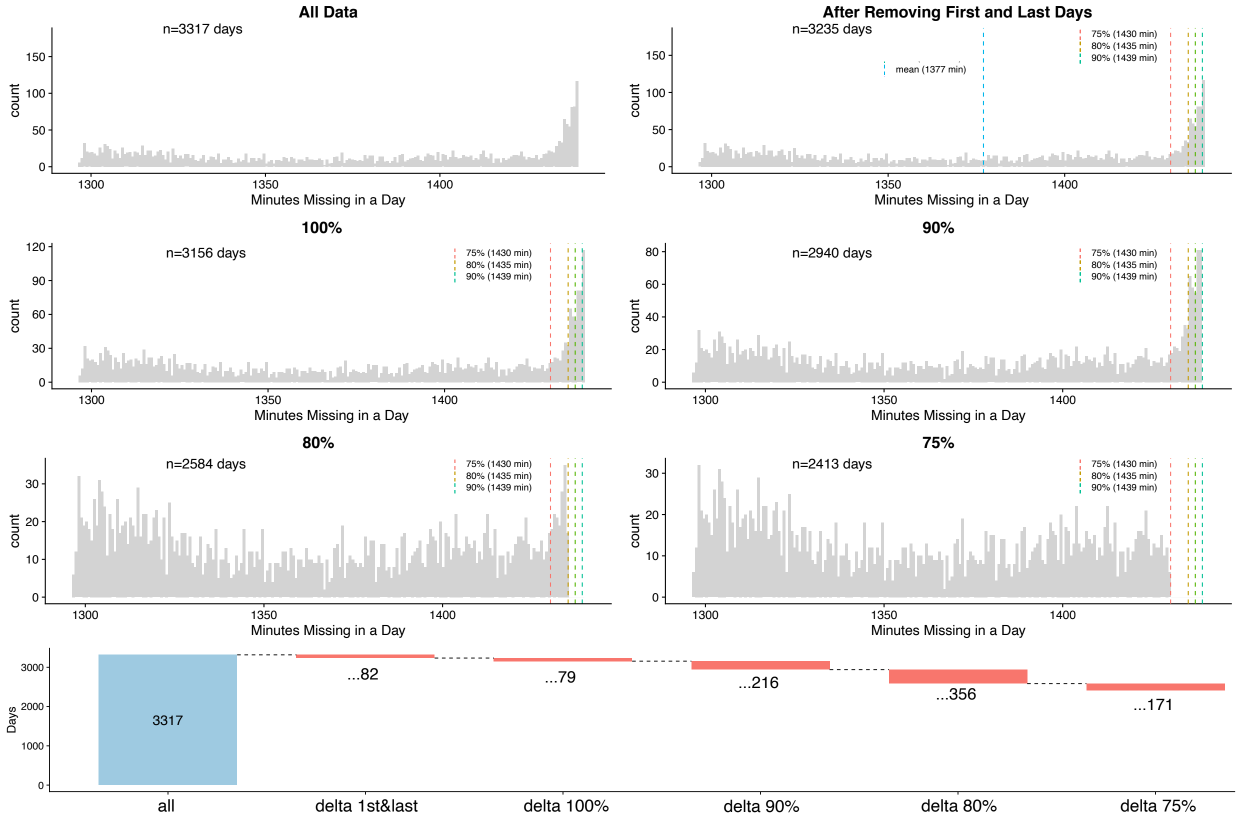

**Supplementary Figure 1 | Mobility data collection and thresholds for missing data.** In total, 3,317 days of recording across all participants were obtained (mean (s.d.) = 77 (26) days, range 14–132 days). After removing the first and last days of each participant's study period when only partial GPS data were recorded (n=82) and days that contained no data (n=79), the remaining data available for analysis contained 3,156 days. To assess the results' sensitivity to data quality, data were then thresholded at three levels 90, 80, and 75 percentiles of missingness. In this case, 100% indicates that a given day was included if any data from that day was available (e.g., missing up to 1,440 minutes, the total number of minutes exist in a day), whereas 75% indicates that days with data missing at the 75$^{th}$ percentile level (missing up to 1,430 minutes).

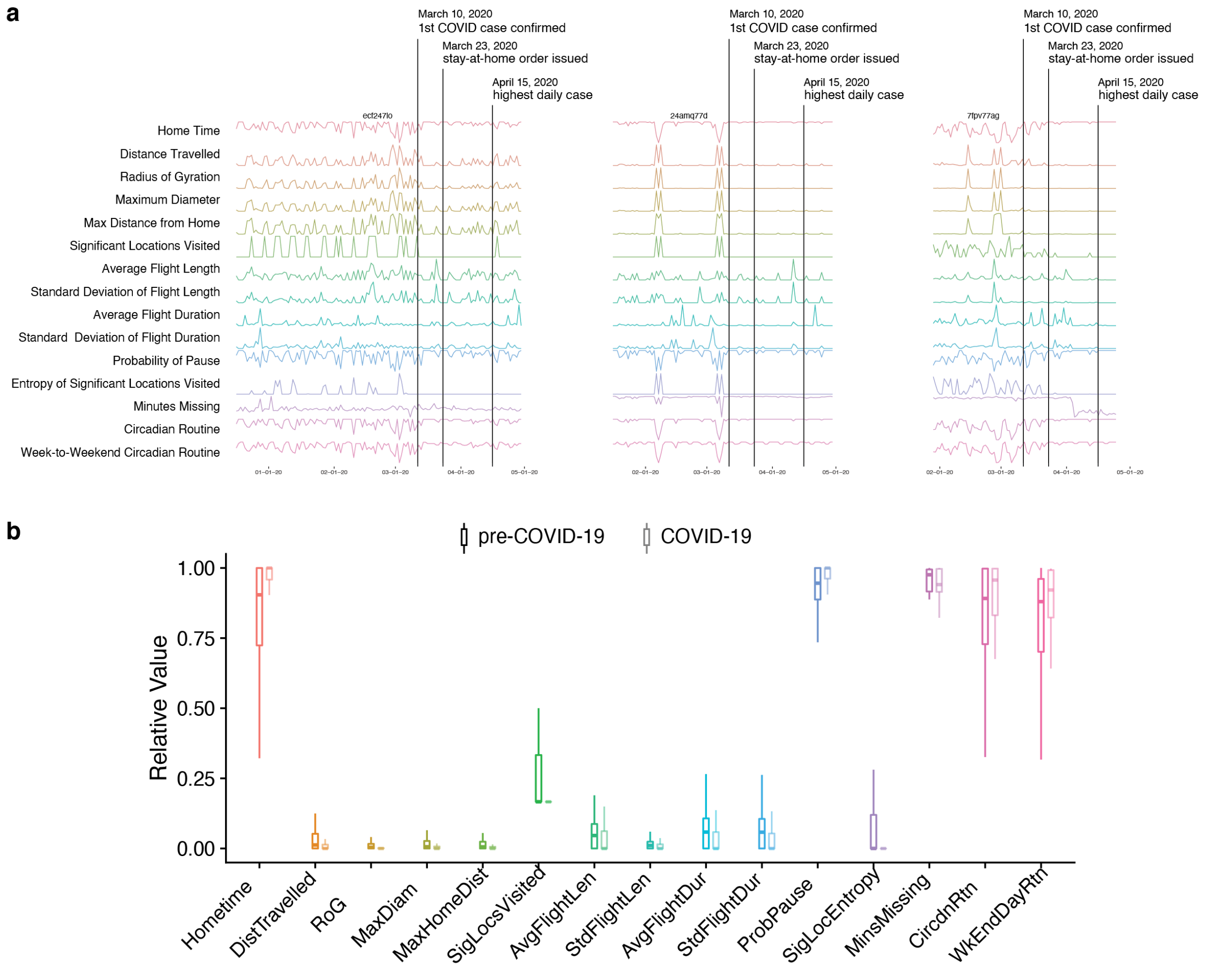

**Supplementary Figure 2 | Dramatic shift in mobility metrics during COVID-19. a)** Three participants still had active mobile phone data collection when COVID-19 hit the Philadelphia area during the end of the study period. Vertical lines indicate milestone dates: 1st confirmed case in Philadelphia (March 10, 2020), stay-at-home order issued by the mayor (March 23, 2020), and the highest daily case in the first wave (April 15, 2020). Visually, mobility metrics across the board exhibited shift since the onset of the pandemic, which we quantified in the bottom panel. **b**) As expected, participants stayed at home longer, travelled less distance, visited fewer locations, took on a more stable circadian routine. However dramatic, data points collected during COVID-19 represented only 1.1% of all data in the current study. Exclusion of these data points did not significantly change the results.

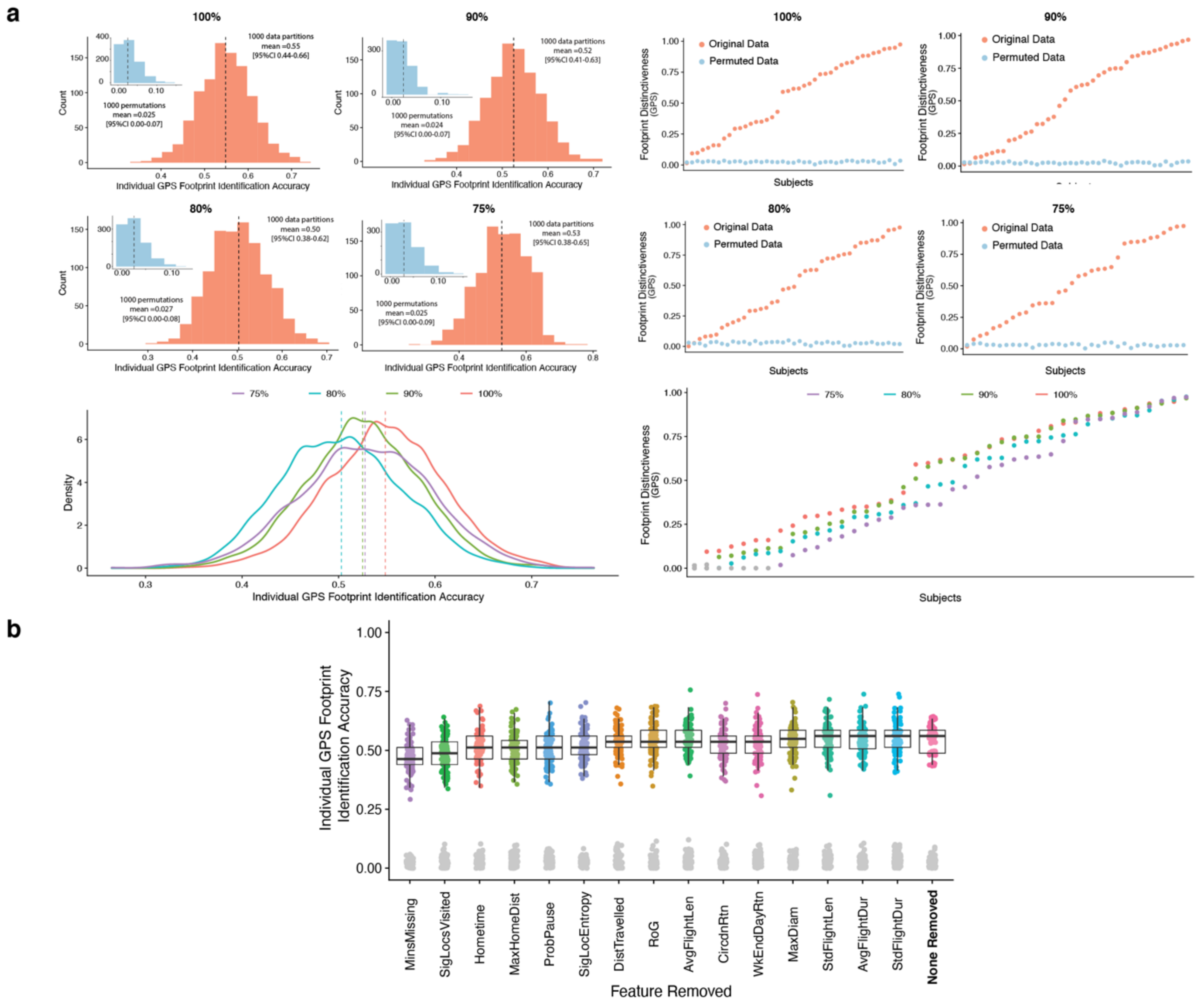

**Supplementary Figure 3 | Convergent results when different thresholds for data missingness are applied or individual mobility features are removed. a**) Histograms of GPS-based identification accuracy as well as subject-level footprint distinctiveness were stable across four inclusion thresholds for data missingness (also see **Supplementary Figure 1**). **b**) Individual identification accuracy was not significantly changed when any of the GPS mobility features was removed from the covariance matrix (“footprint”). Permutation results were shown in grey dots, which demonstrated individual identification were statistically significant regardless which feature was removed.

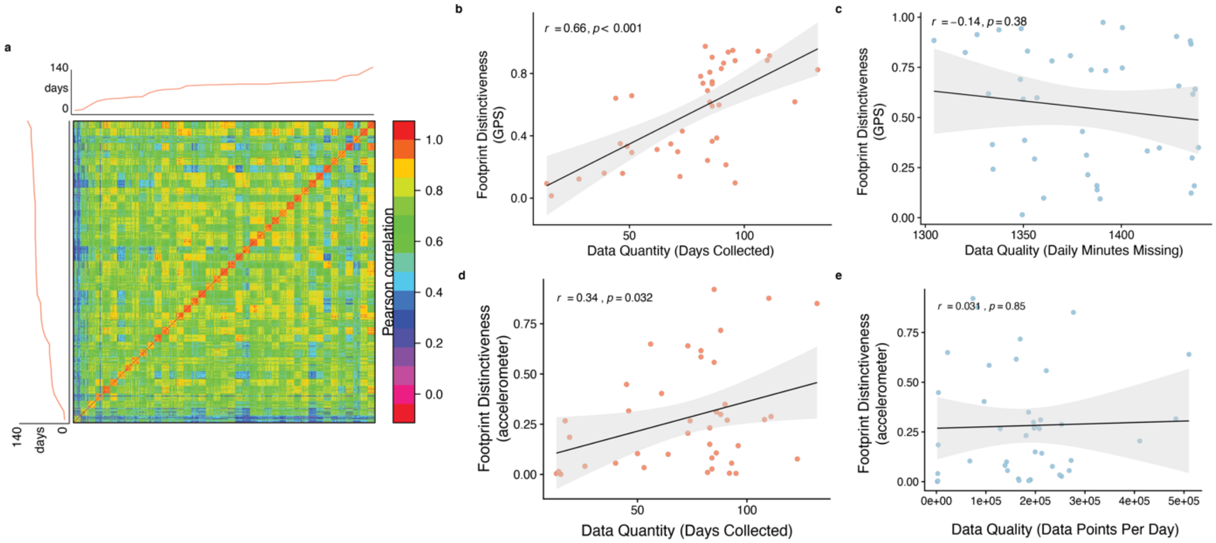

**Supplementary Figure 4 | Individual footprint distinctiveness is driven by data quantity but not data quality. a)** We constructed a subject footprint similarity matrix by grouping together multiple random reference and target partitions from the same subject. The visual contrast between the on and off diagonal block structures represented higher within-subject footprint correlations than between-subject. We observed that when subjects were ranked in order of number of days of geolocation data recorded (red linear plots next to the x and y axes), the prominence of the on-diagonal block structures, or the within-subject similarity, also increased accordingly. Note, grouping was only for visualization and not used in any statistical analysis or individual identification. **b**) We found that data quantity was correlated with footprint distinctiveness ($r = 0.66$, $p < 0.001$). **c**) On the other hand, we found no evidence that data quality (operationalized as the number of minutes missing in daily GPS data) was related to footprint distinctiveness ($p = 0.38$). **d, e**) Similar results were found for accelerometer data. Specifically, data quantity was significantly associated with accelerometer-based footprint distinctiveness ($r = 0.34$, $p < 0.05$), but data quality was not ($p = 0.85$).

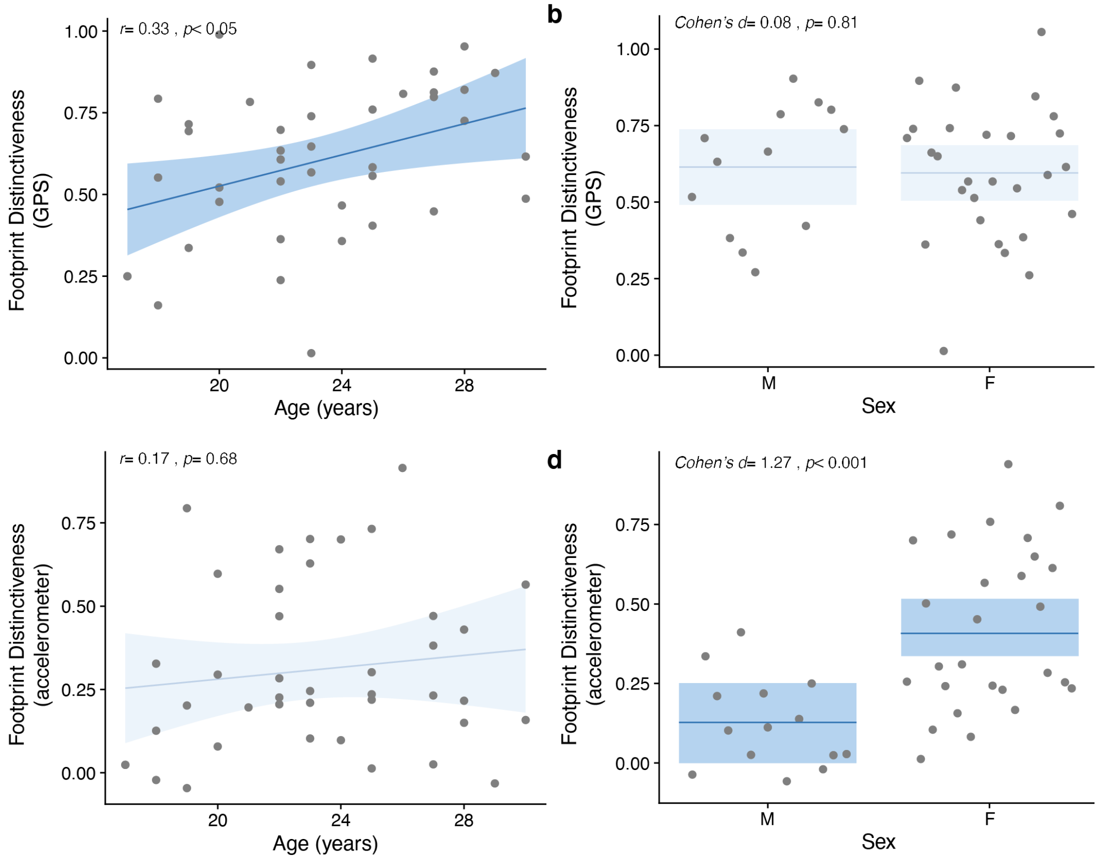

**Supplementary Figure 5 | Developmental effects and sex differences in footprint distinctiveness. a)** While controlling for data quantity, geolocation-based footprint became more distinct with age (partial $r = 0.33$, $p < 0.05$), **b**) but it did not differ between sexes (*Cohen's d* = 0.08, $p = 0.81$). **c**) In contrast, accelerometer data did not exhibit any age effects (partial $r = 0.17$, $p = 0.68$), but females had significantly more distinct accelerometer-based footprint than males (*Cohen'd* = 1.27, $p < 0.001$).

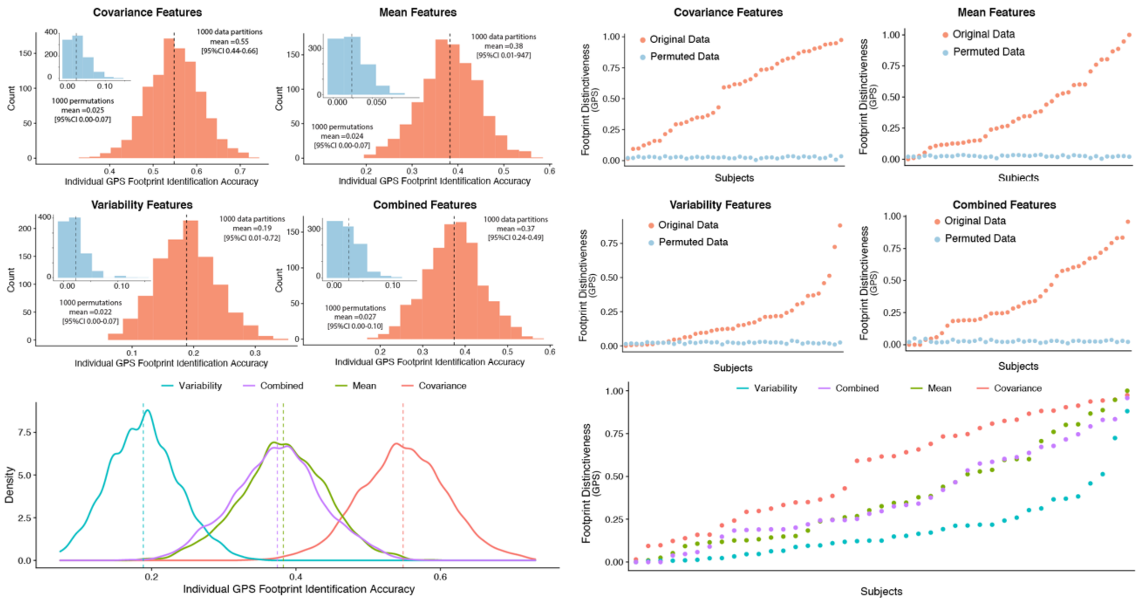

**Supplementary Figure 6 | Identifying individuals using alternative footprint features.** While our main analyses used the covariance of mobility features to identify individuals, we also evaluated alternative feature types. Specifically, we repeated the geolocation-based footprinting analyses using the daily mean value of each feature, the daily variability of each feature (operationalized as the root mean square of the successive differences), and a combined feature set including mean, variability, and covariance features. While all feature sets yielded significant individual identification accuracy compared to the null distribution, covariance features out-performed the alternative features by a wide margin (covariance: 54.8%, mean: 38.2%, variability: 18.9%, combined: 37.4%). This data suggests that covariance features best encoded individual-specific mobility footprints.
